# Repeated hypoglycemia blunts the responsiveness of glucose-inhibited GHRH neurons by remodeling neural inputs and disrupting mitochondrial structure and function

**DOI:** 10.1101/788869

**Authors:** M Bayne, A Alvarsson, K Devarakonda, R Li, M Jimenez-Gonzalez, K. Conner, M Varghese, M N Serasinghe, J E Chipuk, P R Hof, S A Stanley

## Abstract

Hypoglycemia is a frequent complication of diabetes, limiting therapy and increasing morbidity and mortality. With recurrent hypoglycemia, the counter-regulatory response (CRR) to decreased blood glucose is blunted, resulting in hypoglycemia unawareness. The mechanisms leading to these blunted effects remain incompletely understood. Here, we identify, with in situ hybridization, immunohistochemistry and the tissue clearing capability of iDisco, that GHRH neurons represent a unique population of arcuate nucleus neurons activated by glucose deprivation *in vivo*. Repeated glucose deprivation reduces GHRH neuron activation and remodels excitatory and inhibitory inputs to GHRH neurons. We show low glucose sensing is coupled to GHRH neuron depolarization, decreased ATP production and mitochondrial fusion. Repeated hypoglycemia attenuates these responses during low glucose. By maintaining mitochondrial length with the small molecule, mdivi-1, we preserved hypoglycemia sensitivity *in vitro* and *in vivo*. Our findings present possible mechanisms for the blunting of the CRR, broaden significantly our understanding of the structure of GHRH neurons and for the fist time, propose that mitochondrial dynamics play an important role in hypoglycemia unawareness. We conclude that interventions targeting mitochondrial fission in GHRH neurons may offer a new pathway to prevent hypoglycemia unawareness in diabetic patients.

## Introduction

Hypoglycemia is rare in healthy individuals but a frequent occurrence for people with diabetes. Intensive blood glucose control (1) ameliorates neurological and microvascular complications of diabetes but poses a major risk of hypoglycemia. In a prospective study, over 80% of individuals with type 1 diabetes (T1D) and almost half of those with type 2 diabetes experience at least one episode of severe hypoglycemia each month (2). In a recent study using continuous glucose monitoring, individuals with T1D were hypoglycemic (<60 mg/dl) for over 6% of each day (3). Repeated episodes of hypoglycemia blunt the release of counter-regulatory hormones and diminish the physiological responses to subsequent hypoglycemia (4). In turn, hypoglycemia unawareness increases the risk of severe hypoglycemia by up to 20 fold and significantly increases morbidity and mortality (5). The real and perceived risks of hypoglycemia limit effective therapy (6) and reduce well-being in many individuals with diabetes (7).

Under normal circumstances, plasma glucose levels are maintained in a narrow range. Hormones, neurotransmitters and behavior all contribute to glucose homeostasis. In healthy individuals, hypoglycemia rapidly triggers sequential components of the counter-regulatory response (CRR) to restore blood glucose (8). These responses occur via direct actions of low glucose on peripheral organs such as the pancreas and via central nervous system (CNS) actions to regulate peripheral organs through direct innervation and circulating hormones. These include altered pancreatic hormone release, sympathetic activation, increased pituitary (growth hormone (GH) and adrenocorticotrophin hormone (ACTH)) and adrenal (corticosterone/cortisol) hormone release, and behavioral responses such as feeding.

Peripheral and central glucose sensors detect and respond to changes in plasma glucose to play a role in the CRR (9). In the CNS, specialized glucose-sensing neurons use glucose as a signal and can be broadly defined as glucose-excited (GE) or glucose-inhibited (GI) neurons that are activated by high or low glucose respectively. Glucose-sensing neurons are found in many brain regions including hypothalamic nuclei such as the arcuate nucleus (ARC). There is strong evidence for a role for ARC neurons in the response to hypoglycemia. The ARC includes both GE and GI neurons, its neurons express the putative glucose sensors such as glucokinase (10), glucose transporter type 2 (11) and ATP-sensitive potassium channels (12), and are activated by hypoglycemia *in vivo*. Mitochondrial function and structure also contribute to glucose sensing in ARC neurons (13). Disrupting mitochondrial fission or fusion in either ARC GE neurons impairs glucose sensing and leads to systemic metabolic abnormalities (14). ARC neurons are synaptically connected to metabolically active organs such as the pancreas, liver and adipose tissue (15). In addition, ARC neurons play central roles in feeding, a critical behavioral response to hypoglycemia.

A number of mechanisms have been proposed to contribute to hypoglycemia unawareness. These include altered glucose transport into the CNS (16), use of alternate fuels such as lactate (17) or glycogen (18), CNS actions of counter-regulatory hormones such as glucocorticoids (19), modified neurotransmitter release (20) and shifted glucose responses in glucose-sensing neurons (21). However, most brain regions are heterogeneous, comprised of glucose-sensing and non-sensing populations or a combination of GE and GI neurons. Such heterogeneity may mask important adaptations in neural subpopulations.

We have previously shown that ARC growth hormone releasing hormone neurons (GHRH) express glucokinase and are relatively homogeneous as over 80% of GHRH neurons are inhibited by glucose in *ex vivo* electrophysiological studies (10). Therefore, in this current study, we sought to test the hypothesis that repeated glucose deprivation results in structural and functional adaptation of glucose inhibited GHRH neurons leading to inadequate activation with low glucose.

We identified GHRH neurons as a distinct population of ARC GABAergic neurons with polysynaptic inputs to the pancreas that are activated by glucose deprivation *in vivo*. We modeled hypoglycemia insensitivity with repeated glucose deprivation *in vivo* and demonstrated blunted GHRH neuron activation associated with remodeling of excitatory and inhibitory inputs and microglial activation. Further, we generated an *in vitro* model of hypoglycemia insensitivity. *In vitro*, acute glucose deprivation of GHRH neurons elevated intracellular calcium, depolarized neurons, and regulated mitochondrial structure and function. However, repeated glucose deprivation leading to hypoglycemia insensitivity, partially restored ATP production and prevented mitochondrial elongation. Blocking mitochondrial fission with the small molecule, mdivi1, prevented the effects of repeated glucose deprivation and maintained hypoglycemia sensitivity *in vitro* and *in vivo*.

These data suggest glucose-inhibited GHRH neurons may contribute to several components of the CRR while repeated glucose deprivation results in structural and functional remodeling at both circuit and cellular levels that blunt GHRH neuron responses to hypoglycemia. Maintaining mitochondrial fusion preserved activation with low glucose despite repeated glucose deprivation, suggesting interventions that targeting mitochondrial structure and/or function may prevent hypoglycemia unawareness.

## Results

### GHRH neurons are multipolar neurons with sparse dendritic processes

We first characterized the distribution and morphology of GHRH neurons *in vivo*. We examined the number and distribution of GFP-positive neurons in iDISCO+ (22) cleared brains from previously validated transgenic mice expressing GFP in GHRH neurons (GHRH-GFP mice) (23) (Fig 1A). GHRH neurons represent a population of 354 ± 10 neurons per brain with expression primarily in the ARC (82.9 ± 4.3%) and a minor population in the periventricular nucleus (14.3 ± 4.3%). Alexa-filled ARC GHRH-GFP neurons were uniformly multipolar, with three or more dendrites (Fig 1B). GHRH neurons have sparse dendritic spines (0.1 ± 0.02 spines/µm) of which the majority were thin spines or filopodia (65.9%), which can be rapidly remodeled. Stubby (23.4%) and mushroom (10.6%) spines were also present. Spine density did not show any difference with distance from the soma for any spine type (Fig 1C-D).

**FIGURE 1:**
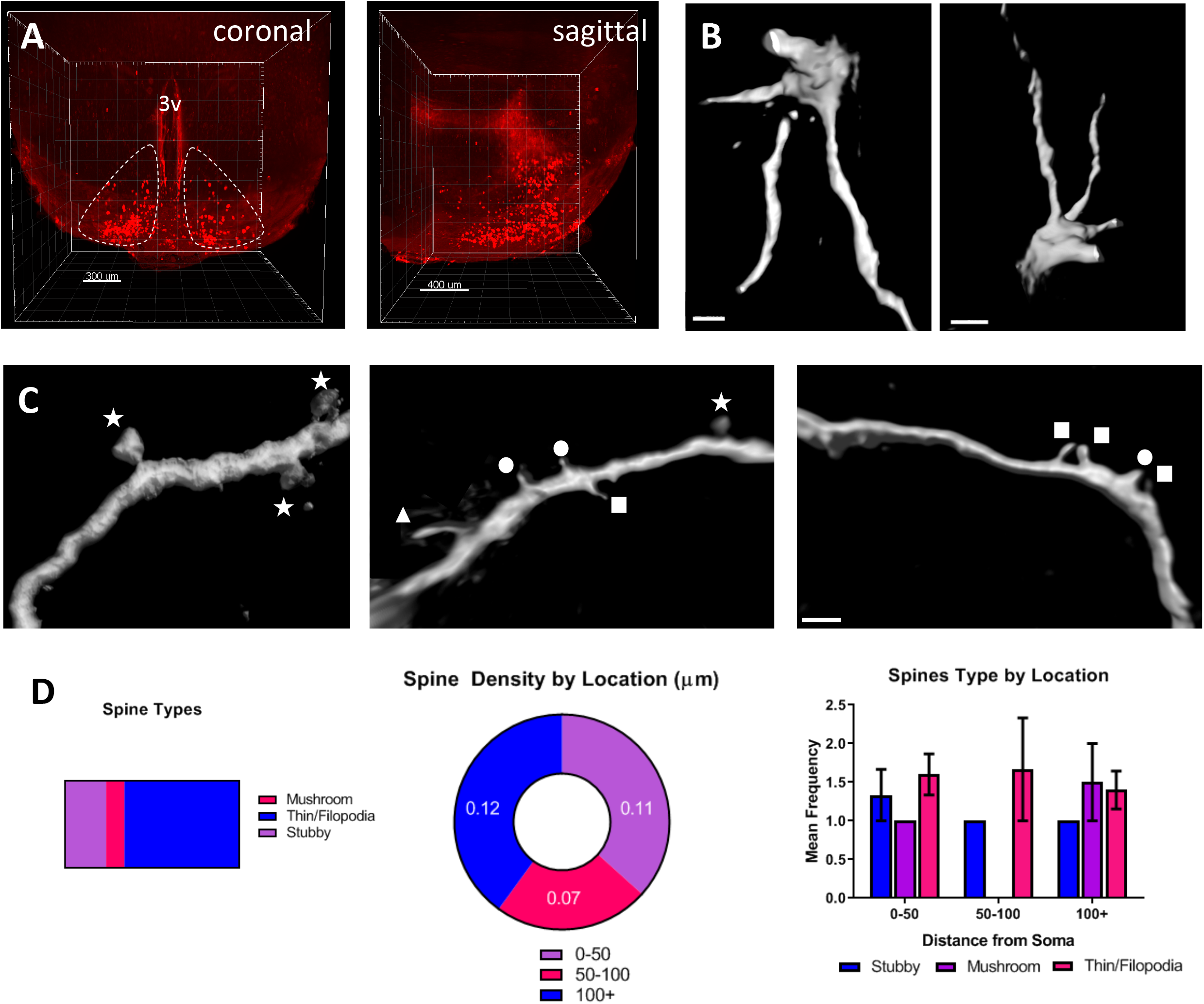
GHRH neurons are multipolar neurons with sparse dendritic processes. A) Light sheet images of brain from GHRH-GFP mice with optical clearing by iDISCO showing distribution of GFP+ GHRH neurons in coronal and sagittal planes. GFP positive neurons are pseudo-colored in red. B) 3D reconstruction of multipolar GHRH neurons after Alexa-555 filling and confocal imaging. Scale bar = 2 µm left panel, scale bar = 3 µm right panel. C) Maximum intensity projections of GHRH neurons demonstrating dendritic spine types: thin 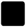 filopodia 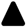, mushroom 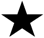 and stubby 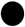. Scale bar = 1 µm for all panels D) Proportion of dendritic spine types on filled GHRH neurons (left). Distribution of all dendritic spines with distance from the soma (center) and distribution of specific spine types with distance from the soma (right) (n = 47).

### GHRH neurons are non-AGRP, non-POMC GABAergic neurons synaptically connected to the pancreas

The ARC contains several neural populations with distinct physiological roles, particularly in relation to feeding, which is a crucial response to hypoglycemia. To determine the neurochemical phenotype of GHRH neurons, fluorescent *in situ* hybridization (FISH) and immunohistochemical (IHC) labeling approaches were used. *In situ* hybridization for *VGLUT2*, *GAD2*, and *GHRH* revealed GHRH neurons were primarily GABAergic (Fig 2A). In our analysis, all *GHRH* neurons expressed *GAD2* mRNA and there was no detectable colocalization of *GHRH* and *VGLUT2*. Neuropeptide expression was determined by IHC in brain slices from GHRH-GFP mice after colchicine injection. Only 0.8 ± 0.5% of GHRH neurons colocalize with neuropeptide Y (NPY)-expressing neurons and there was no detectable overlap between GHRH neurons and neurons expressing agouti related peptide (AGRP) or between GHRH and neurons expressing pro-opiomelanocortin (POMC) (Fig 2B). These data are consistent with GHRH neurons being a distinct population of ARC GABAergic, non-AGRP, non-POMC neurons.

**FIGURE 2:**
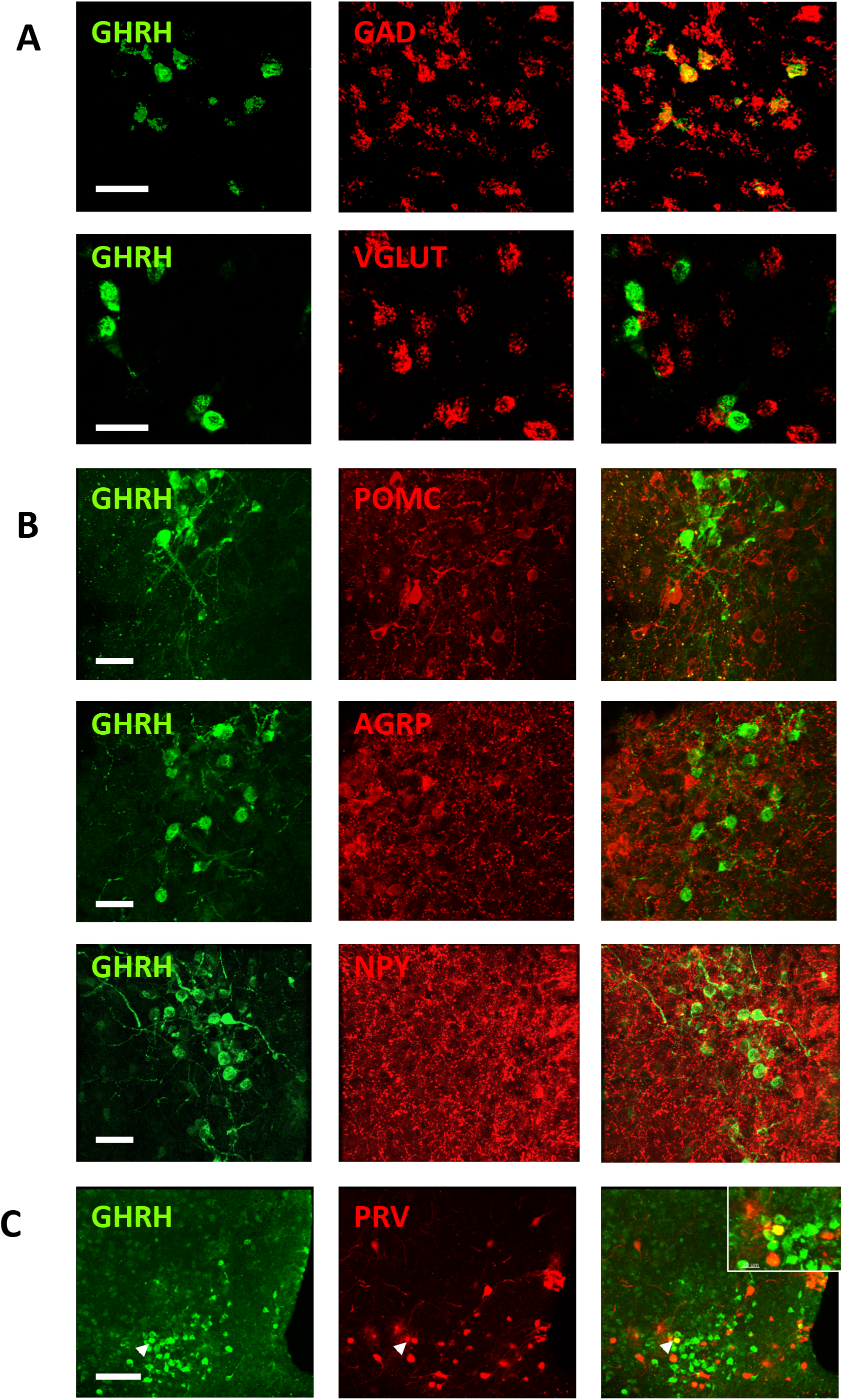
Arcuate GHRH neurons are non-Agrp, non-POMC GABAergic neurons synaptically connected to the pancreas. A) Confocal analyses of fluorescent *in situ* hybridization for *GHRH* and *GAD* (upper) or *VGLUT* (lower panel) in ARC (n = 4/marker). Scale bar: 30 µm for all panels B) Confocal analyses of immunohistochemistry for GFP and POMC (upper), AGRP (middle) or NPY (lower panel) in ARC of GHRH-GFP mice (n = 5/marker). Scale bar: 30 µm for all panels C) Confocal analyses of immunohistochemistry for GFP and PRV-RFP in the ARC after intrapancreatic injection of PRV-RFP in GHRH-GFP mice. Inset: Overlap between GFP and RFP immunolabeling (n = 5). Scale bar: 80 µm for all panels

Next, we investigated if GHRH neurons were synaptically connected to peripheral organs involved in CRR. In GHRH-GFP mice injected with the viral retrograde tracer, pseudorabies virus (PRV-RFP), a subpopulation of GHRH neurons colocalized with RFP expression after PRV-RFP injection into the pancreas (1.3 ± 0.5% of GHRH-GFP cells, 5.1 ± 1.9% of ARC PRV-RFP-immunoreactive neurons) (Fig 2C). There was no overlap between GHRH-GFP-positive neurons and RFP-positive neurons after PRV injection into liver, muscle, epididymal white fat or adrenal (data not shown). These data demonstrate that a subpopulation of GHRH neurons forms a polysynaptic pathway to innervate the pancreas, and as such may be involved in direct neuronal regulation of pancreatic function.

### Repeated glucose deprivation with peripheral 2-deoxy-D-glucose blunts the CRR

Systemic glucose deprivation was achieved by intraperitoneal (i.p.) administration of 2-deoxy-D-glucose (2DG) (24), a non-metabolizable glucose analog that mimics the physiological response to hypoglycemia (Fig 3A). We chose to use 2DG rather than insulin for glucose deprivation because insulin receptor expression has been reported in ARC neurons and so insulin administration could result in neural activation via receptor binding in the ARC, independent of glucose deprivation. 2DG injection (400 mg/kg i.p.) resulted in a counter-regulatory increase in blood glucose in 6-hour fasted C57BL6 mice reaching a maximum at 90 mins. Based on this time-course, all other measurements were made at the 90 min time-point. Repeated glucose deprivation (2DG injection i.p. daily for 5 days) recapitulated the blunted CRR seen with repeated hypoglycemia. Peak blood glucose 90 minutes after 2DG was significantly reduced compared to acute glucose deprivation (Fig 3B). A single episode of glucose deprivation with 2DG significantly increased plasma glucagon and this effect was blunted by repeated glucose deprivation to 25% of the glucagon response with acute glucose deprivation (Fig 3C). Similarly, plasma corticosterone was significantly increased by glucose deprivation compared to vehicle treated control and reduced by 30% with repeated glucose deprivation (Fig 3D). In contrast to humans where hypoglycemia acutely increases plasma GH, there was no significant change in plasma GH with acute or repeated glucose deprivation compared to GH in normoglycemic mice but GH was significantly higher after repeated glucose deprivation than after acute glucose deprivation (Fig 3E).

**FIGURE 3:**
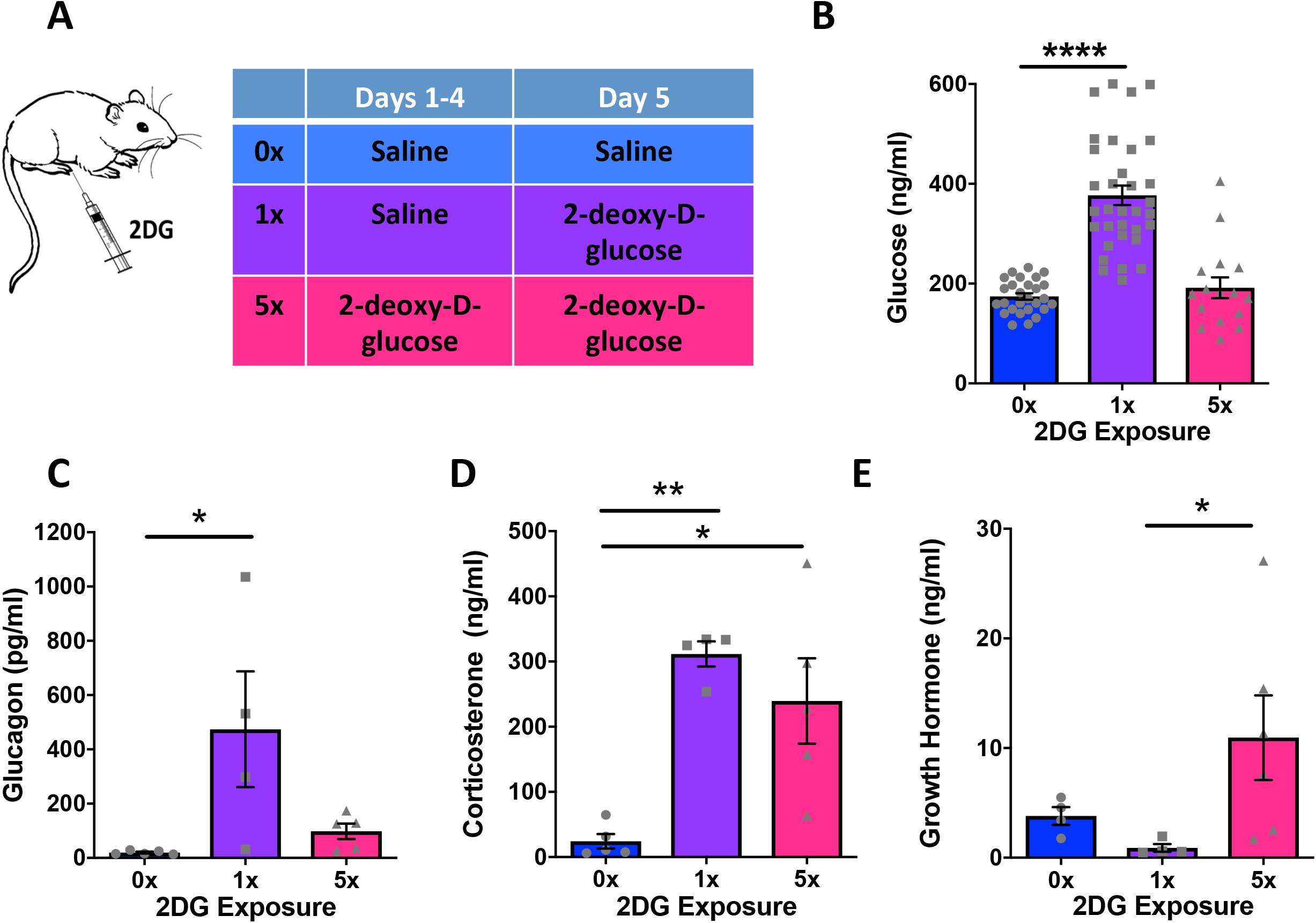
Repeated glucose deprivation with 2 deoxyglucose blunts the counter-regulatory response. A) Schema of experimental protocol for repeated glucose deprivation with intraperitoneal administration of 2-deoxyglucose (2DG) *in vivo*. B – E) Analysis of blood glucose (B) (**** p < 0.0001, χ^2^_(2)_ = 47.4, Kruskal-Wallis ANOVA with Dunn’s multiple comparison test, n = 16 – 33/treatment group), plasma glucagon (C) (* p = 0.03, F_(2,11)_ = 4.91, one way ANOVA with Tukey’s multiple comparison test, n = 4-5/group), plasma corticosterone (D) (* p= 0.01, ** p = 0.001, F_(2,11)_ = 12.33, ANOVA with Tukey’s multiple comparison test, n = 4-5/treatment group) and plasma growth hormone (E) (* p = 0.02, χ^2^_(2)_ = 7.12, Kruskal-Wallis ANOVA with Dunn’s multiple comparison test, n = 4-6/group) after vehicle (0x), single (1x) or repeated (5x) i.p. 2DG administration.

### GHRH neuron activation by acute glucose deprivation is impaired by repeated glucose deprivation

Previous electrophysiological recordings in *ex vivo* brain slices demonstrated that GHRH neurons are activated by low glucose. Thus, we next wanted to confirm GHRH neuron activation by glucose deprivation *in vivo* and test the hypothesis that this activation is blunted by prior glucose deprivation. Quantitative analysis of the early immediate gene *c-fos* across the whole ARC using FISH showed no significant effect of acute or repeated glucose deprivation compared to fasted, vehicle-treated controls (Fig 4A). Combined FISH for *GHRH* and *c-fos* revealed selective activation of GHRH neurons by glucose deprivation. Acute glucose deprivation increased c-fos/GHRH-expressing neurons by 10 fold and this effect was abolished by repeated glucose deprivation (Fig 4B). Next, we assessed whether cell activation was accompanied by transcriptional changes in GHRH neurons. Acute glucose deprivation doubled *GHRH* mRNA probe density compared to vehicle treatment and this increase was absent with repeated glucose deprivation (Fig 4C). In addition, acute hypoglycemia increased the density of the putative glucose sensor, glucokinase (GK) in GHRH neurons by three fold (Fig 4D). These data are consistent with GHRH neuron activation by glucose deprivation *in vivo* and suggest rapid upregulation of *GHRH* and *GK* expression in response to low glucose. These effects are lost with repeated glucose deprivation.

**FIGURE 4:**
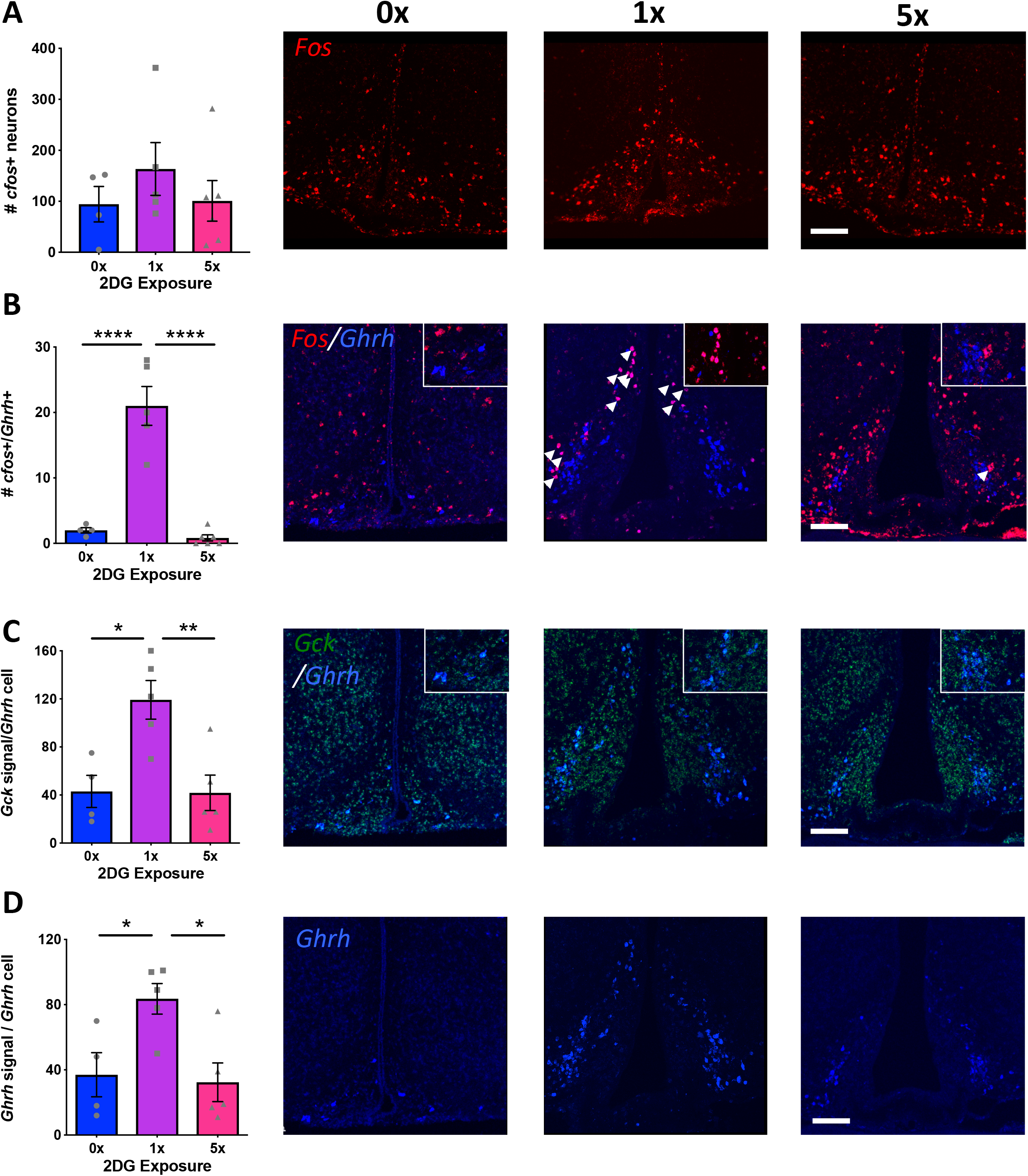
GHRH neuron activation by acute glucose deprivation is impaired by repeated glucose deprivation. A) Quantification and confocal analyses of fluorescent *in situ* hybridization for *cfos* in the ARC after vehicle (0x), single (1x) or repeated (5x) i.p. 2DG administration. n = 4-6/group. B) Quantification of *cfos*-positive and *Ghrh* positive cells and confocal analyses of fluorescent *in situ* hybridization for *cfos* and *GHRH* in the ARC after vehicle (0x), single (1x) or repeated (5x) i.p. 2DG administration, **** p < 0.0001 vs. 0x and vs. 5x F _(2, 12)_ = 42.1, one way ANOVA with Tukey’s multiple comparison test, n = 4 – 6/group. C) Quantification of glucokinase expression in *Ghrh* positive cells and confocal analyses of fluorescent *in situ* hybridization for *Gck* and *Ghrh* in the ARC after vehicle (0x), single (1x) or repeated (5x) i.p. 2DG administration. * p = 0.013 vs. 0x and ** p = 0.008 vs. 5x, F_(2, 11)_ = 8.94, one way ANOVA with Tukey’s multiple comparison test, n = 4 – 5/group. D) Quantification of *Ghrh* expression and confocal analyses of fluorescent *in situ* hybridization for *Ghrh* in the ARC after vehicle (0x), single (1x) or repeated (5x) i.p. 2DG administration. * p = 0.04 vs. 0x and ** p = 0.02 vs. 5x, F_(2, 11)_ = 6.34, one way ANOVA with Tukey’s multiple comparison test, n = 4 – 5/group. Scale bar: 30 µm for all panels. Each dot represents results from individual animals and data are displayed as mean ± SEM.

### Repeated glucose deprivation results in morphological changes in GHRH neurons and activated microglia

Low glucose activation of GHRH neurons could be a direct effect or secondary to alterations in the balance of excitatory and inhibitory inputs onto GHRH neurons. Dendritic spines are the primary sites of excitatory glutamatergic synapses in mammalian brains and can undergo remodeling (25). Increased excitatory inputs increase spine number and conversely, dendritic spines are lost with reduced excitatory inputs and synaptic long-term depression. Therefore, we investigated the effects of acute and repeated glucose deprivation on dendritic spines in filled GHRH neurons (Fig 5A and B). Acute glucose deprivation with 2DG had no significant effect on GHRH dendritic spine number compared to those from fasted, vehicle-treated mice. However, in mice treated with repeated episodes of glucose deprivation, we were unable to detect any dendritic spines on GHRH neurons. These findings are in keeping with direct activation of GHRH neurons by acute glucose deprivation but significant loss of excitatory inputs with repeated glucose deprivation.

**FIGURE 5:**
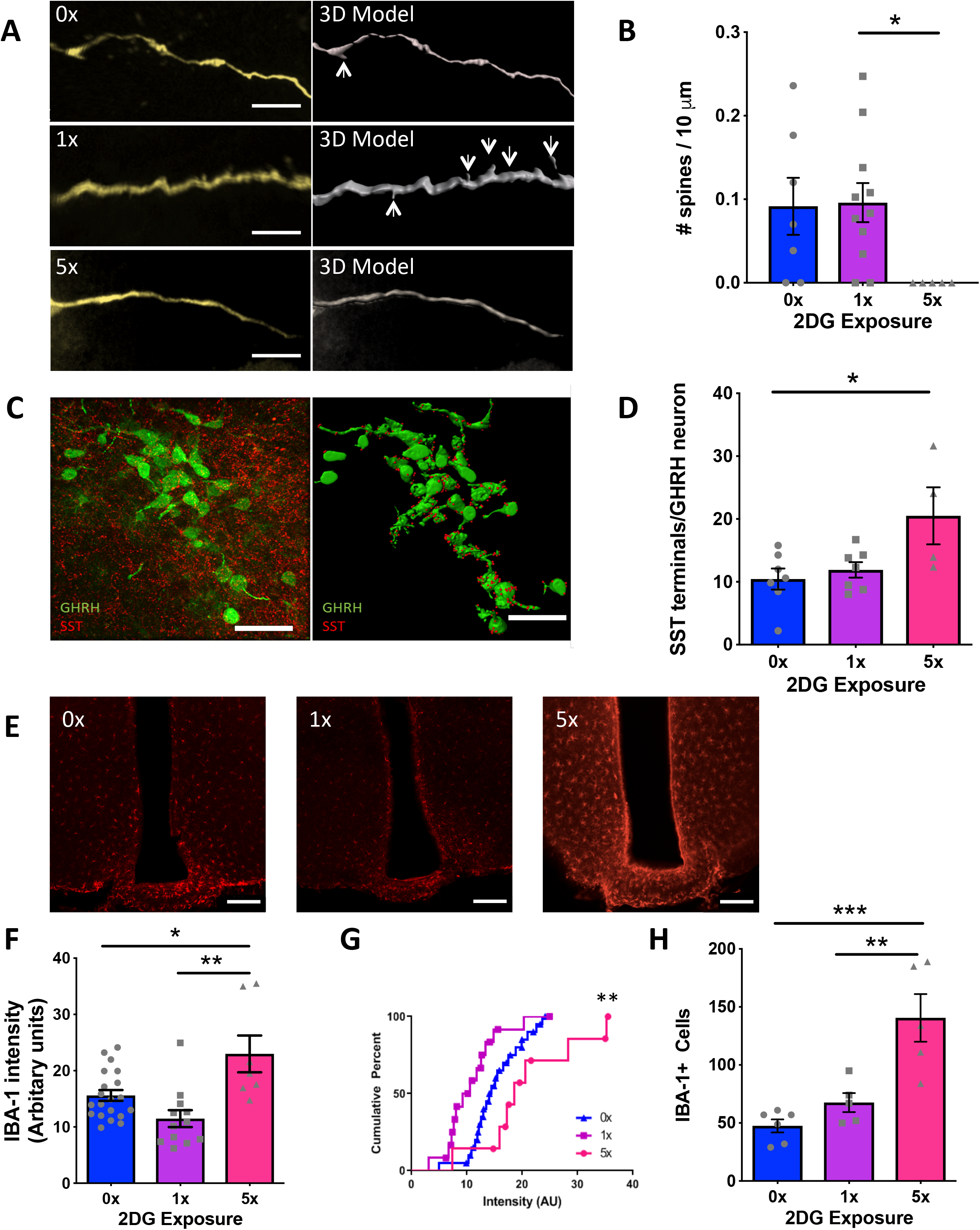
Repeated glucose deprivation disrupts inputs onto GHRH neurons and activates microglia. A) Confocal analyses (left) and 3D reconstruction (right) of dendritic spines on LY-filled ARC GHRH-GFP neurons after vehicle (0x), single (1x) or repeated (5x) i.p. 2DG administration. Scale bar: 5 µm for all panels. B) Quantification of dendritic spines on filled ARC GHRH-GFP neurons after vehicle (0x), single (1x) or repeated (5x) i.p. 2DG administration. * p = 0.03 1x vs. 5x, χ^2^_(2)_ = 7.305, Kruskal-Wallis test with Dunn’s multiple comparison test, n = 5 −11/group. C) Confocal analysis (left) and 3D model (right) of somatostatin-immunoreactive terminals contacting ARC GHRH-GFP neurons. Scale bar: 30 µm D) Quantification of somatostatin-immunoreactive terminals contacting ARC GHRH-GFP neurons after vehicle (0x), single (1x) or repeated (5x) i.p. 2DG administration. * p = 0.02 0x vs. 5x, F_(2, 15)_ = 4.87, one-way ANOVA with Tukey’s multiple comparison test, n = 4 −7/group. E) Confocal analyses of ARC IBA1-positive microglia after vehicle (0x), single (1x) or repeated (5x) i.p. 2DG administration. F) Quantification of IBA-1 intensity by immunohistochemistry in all sections in the ARC after vehicle (0x), single (1x) or repeated (5x) i.p. 2DG administration. * p = 0.01 0x vs. 5x, ** p = 0.0003 1x vs. 5x, F_(2,36)_ = 9.612, one-way ANOVA with Tukey’s multiple comparison test, n = 7 −20/group. G) Cumulative intensity distribution of IBA1 intensity ** p = 0.028 0x vs. 5x, Kolmogorov-Smirnov test n = 7 −20/group. H) Quantification of IBA1-positive cells in the ARC after vehicle (0x), single (1x) or repeated (5x) i.p. 2DG administration. ** p = 0.003, *** p = 0.0004, F_(2,13)_ = 15.35, one-way ANOVA with Tukey’s multiple comparison test, n = 5-6/group. Each dot represents results from individual animals and data are displayed as mean ± SEM.

Next, we measured the density of somatostatin (SST)-immunoreactive terminals, a major inhibitory input, on GHRH neurons. Acute glucose deprivation did not significantly alter the number of SST-containing terminals contacting GHRH neurons but repeated glucose deprivation doubled the number of these terminals in apparent apposition with GHRH neurons (Fig 5C and D). There was no significant effect of acute or repeated glucose deprivation on *c-fos* expression in ARC *Sst*-expressing neurons. These data support the notion that GHRH neurons are directly activated in response to low glucose but suggest that increased inhibitory inputs contribute to loss of GHRH activation with repeated glucose deprivation.

Microglia have been implicated in synaptic pruning in response to physiological insults during development and through life (26), and may play a role in the remodeling of excitatory and inhibitory inputs onto GHRH neurons. To test this hypothesis, we investigated whether acute or repeated glucose deprivation regulated ARC microglia activation using ionized calcium binding adaptor molecule 1 (Iba1) as a marker of activated microglia (Fig 5E). IBA1 expression was not significantly altered by acute glucose deprivation but repeated glucose deprivation increased IBA1 staining intensity by 50% (Fig 5F) and significantly increased the number of IBA1-expressing cells (Fig 5G) as compared to acute glucose deprivation. These data are consistent with an increase in microglial activation with repeated glucose deprivation.

### Repeated exposure of clonal GHRH neurons to low glucose blunts GHRH release

To assess the cell intrinsic responses of GHRH neurons to acute and repeated glucose deprivation, we sought to recapitulate hypoglycemia unawareness with repeated glucose deprivation *in vitro* (1, 3 or 5 episodes) (Fig 6A). We confirmed that the mouse hypothalamic cell line N38 secreted GHRH. GHRH release from N38 cells was increased by 40 – 50% with hypoglycemia (Fig 6B) (single and 3 episodes) but blunted by repeated hypoglycemia (5 episodes). Expression of *GHRH* mRNA was not significantly increased by a single episode of hypoglycemia but repeated hypoglycemia (5 episodes) significantly reduced GHRH expression (Fig 6C). These data suggest repeated exposure of N38 cells to low glucose blunts GHRH release and provides an *in vitro* model for impaired glucose sensing.

**FIGURE 6:**
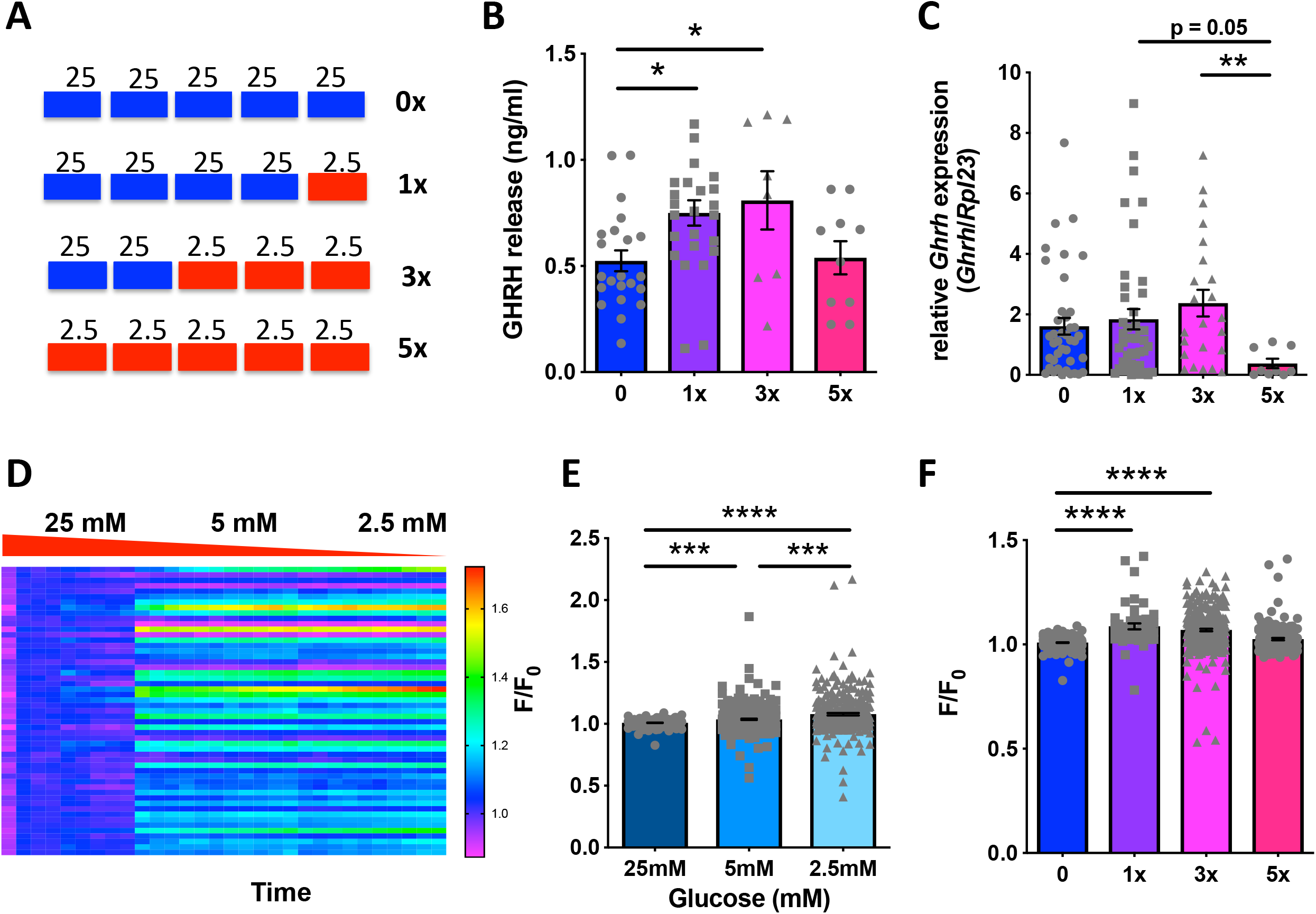
N38 Cells are glucose inhibited and responses are blunted by recurrent glucose deprivation. A) Schema of experimental protocol for repeated glucose deprivation of N38 cells *in vitro* by treatment with media containing 25 mM glucose (standard culture conditions) or 2.5 mM glucose (glucose deprivation). B) Quantification of GHRH release from N38 cells after no (0x), single (1x) or repeated glucose deprivation (3x and 5x; n = 3-4 experiments, in triplicate). * p = 0.04, F_(3, 61)_ = 3.854, one-way ANOVA with Tukey’s multiple comparison test. C) Quantification of *Ghrh* expression in N38 cells after no (0x), single (1x) or repeated glucose deprivation (3x and 5x) (n = 3-4 experiments, in triplicate). ** p = 0.004, χ^2^_(3)_ = 11.61, Kruskal-Wallis test with Dunn’s multiple comparison test. D) Time-resolved calcium responses using calcium indicator Fluo-4 (F/F_0_, color scale) of 53 N38 cells without previous glucose deprivation (1 cell per row) with 25 mM, 5 mM and 2.5 mM glucose treatment. E) Quantification of peak fluorescence (F/F_0_) with 25 mM, 5 mM and 2.5 mM glucose treatment in N38 cells without previous glucose deprivation (4 studies, 51 - 307 cells). *** p = 0.0004, 25 mM vs. 5 mM, *** p = 0.0008, 5 mM vs. 2.5 mM, **** p < 0.0001, 25mM vs. 2.5mM, χ^2^_(2)_ = 56.2, Kruskal-Wallis test with Dunn’s multiple comparison test. F) Quantification of peak fluorescence (F/F_0_) with no (0x), single (1x) or repeated glucose deprivation (3x and 5x) in N38 cells (4 studies, 170-307 cells). **** p < 0.0001, χ^2^_(3)_ = 188.2, Kruskal-Wallis test with Dunn’s multiple comparison test. Each dot represents data from individual cells and data are displayed as mean ± SEM.

### Blunted calcium entry and depolarization of clonal GHRH neurons with repeated low glucose

Because GHRH release is calcium-dependent, we examined the effects of low glucose on intracellular calcium in N38 cells, with and without previous exposure to low glucose. The intensity of the fluorescent calcium indicator, Fluo-4, significantly increased as glucose levels decreased (Fig 6D and E) in keeping with increased intracellular calcium. However, previous episodes of hypoglycemia significantly blunted the calcium response to low glucose such that it was no longer significantly different from baseline (Fig 6F). Next, we assessed the effects of glucose on the voltage response in N38 cells as voltage-gated calcium channels typically mediate neuronal calcium changes. In keeping with our previous electrophysiological findings in GHRH neurons *ex vivo*, N38 cells were depolarized by lowering glucose, as indicated by significantly increased fluorescence intensity of Fluovolt (Fig 7A and B). However, repeated hypoglycemia (5x) reduced baseline fluorescence intensity in keeping with hyperpolarization of N38 cells (Fig 7C). In addition, depolarization in response to low glucose was significantly blunted by previous hypoglycemia (Fig 7D). Together, these data suggest that repeated glucose deprivation hyperpolarizes N38 cells and blunts depolarization in response to low glucose.

**FIGURE 7:**
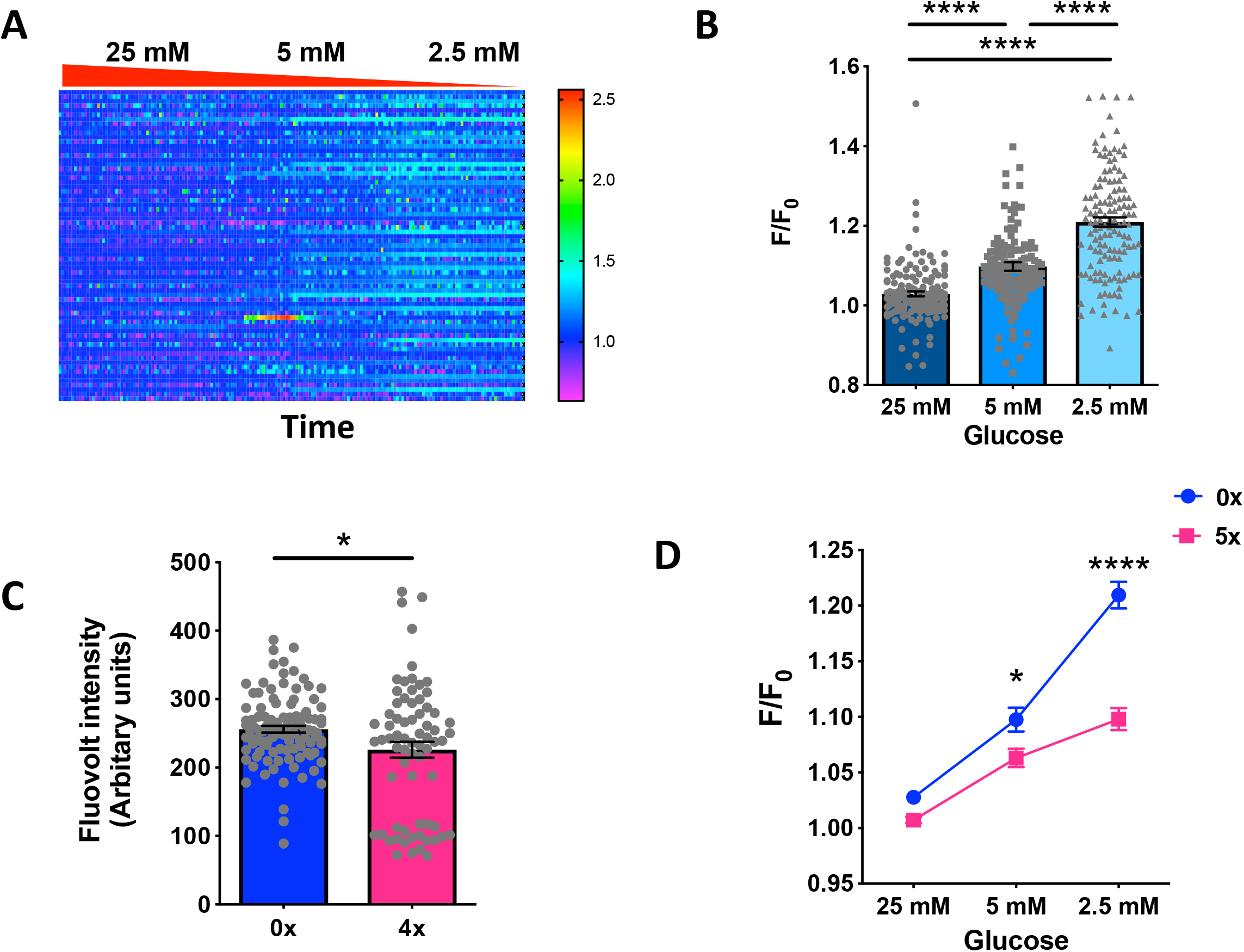
Recurrent glucose deprivation leads to hyperpolarization and impaired depolarization with low glucose. A) Time resolved voltage responses using voltage indicator Fluovolt (F/F_0_, color scale) of 69 N38 cells without previous glucose deprivation (1 cell per row) with 25 mM, 5 mM and 2.5 mM glucose treatment. B) Quantification of peak fluorescence (F/F_0_) with 25 mM, 5 mM and 2.5 mM glucose treatment in N38 cells without previous glucose deprivation (4 studies, 135-144 cells/group). **** p < 0.0001, χ^2^_(2)_ = 153.9, Kruskal-Wallis test with Dunn’s multiple comparison test. C) Quantification of basal fluorescence at 25 mM glucose in N38 cells without (0x) and with (5x) previous glucose deprivation (4 studies, 80 −105 cells). p = 0.02, *t* = 2.371, df = 93.5, unpaired *t*-test with Welch’s correction. D) Quantification of peak fluorescence (F/F_0_) with 25 mM, 5 mM and 2.5 mM glucose treatment in N38 cells without previous glucose deprivation (4 studies, 195 cells). * p = 0.03, **** p < 0.0001, significant effect of previous glucose deprivation, F_(193, 386)_ = 1.915, p < 0.0001, two-way ANOVA with Sidak’s multiple comparisons test. Each dot represents data from individual cells and data are displayed as mean ± S.E.M.

### Repeated glucose deprivation blunts low glucose effects on ATP production and mitochondrial length

Increased reactive oxygen species (ROS) have been proposed to play a critical role in glucose sensing in both ARC GE neurons and in response to insulin-induced hypoglycemia in VMH neurons. Therefore, we measured the effects of acute and repeated low glucose on whole cell ROS in N38 cells. Repeated glucose deprivation significantly decreased whole cell ROS, measured by intensity of the fluorescent probe, Amplite Fluorescent Green (27), compared to acute deprivation (Fig 8A). These findings are consistent with decreased ROS signaling with repeated glucose deprivation.

**FIGURE 8:**
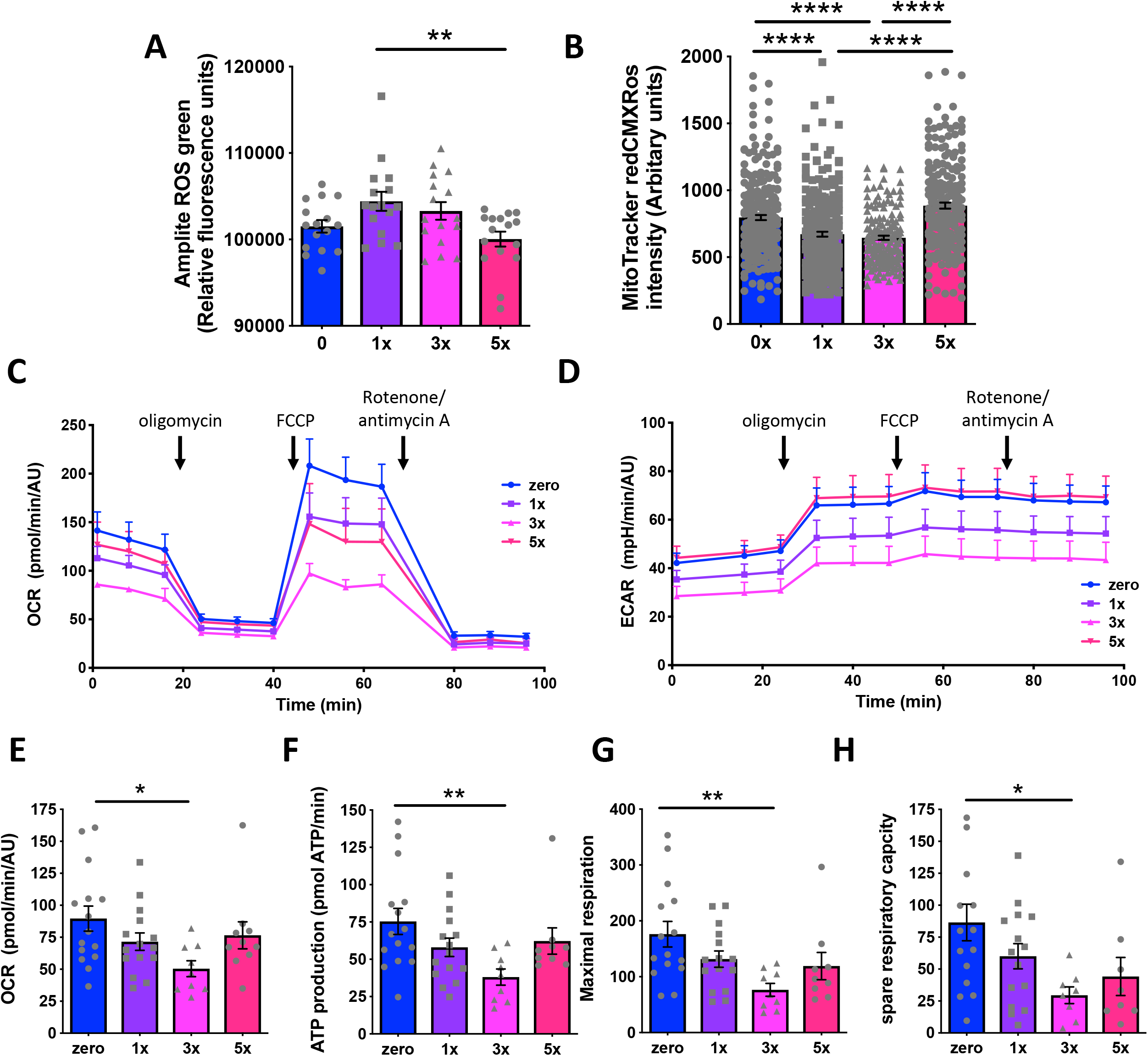
Repeated glucose deprivation blunts the effects of low glucose on mitochondrial function. Quantification in N38 cells after none (0x), single (1x) or repeated glucose deprivation (3x and 5x) A) Amplite ROS green whole cell reactive oxygen species indicator (n = 3-4 experiments, in triplicate). ** p = 0.008 1x vs. 5x, F_(3, 60)_ = 4.246, one-way ANOVA with Tukey’s multiple comparison test. B) Mitotracker Red CMX intensity (n = 3-4 experiments, in triplicate, 160 – 252 cells/group). **** p < 0.0001, χ^2^_(3)_ = 83.5, Kruskal-Wallis test with Dunn’s multiple comparison test. C) Oxygen consumption rate (OCR) (n = 3-4 experiments, in triplicate). D) Extracellular acidification rate (ECAR) (n = 3-4 experiments, in triplicate). E) Basal respiration rate (n = 3-4 experiments, in triplicate). * p = 0.02, χ^2^_(3)_ = 8.665, Kruskal-Wallis test with Dunn’s multiple comparison test. F) ATP production (n = 3-4 experiments, in triplicate). ** p = 0.007, χ^2^_(3)_ = 10.66 Kruskal-Wallis test with Dunn’s multiple comparison test. G) Maximal respiration (n = 3-4 experiments, in triplicate). ** p = 0.001, χ^2^_(3)_ = 12.45, Kruskal-Wallis test with Dunn’s multiple comparison test. H) Spare respiratory capacity (n = 3-4 experiments, in triplicate). * p = 0.04, χ^2^_(3)_ = 9.015, Kruskal-Wallis test with Dunn’s multiple comparison test. Each dot represents data from individual well and data are displayed as mean ± S.E.M.

ATP has also been proposed to be a critical intracellular signal for glucose sensing and to play a role in synaptic morphology, which we demonstrated are both disrupted by repeated glucose deprivation. Mitochondrial function is crucial for both ROS and ATP production. To understand effects of acute and repeated glucose deprivation on mitochondrial function, we first assessed mitochondrial membrane potential. Mitochondrial membrane potential, determined by fluorescence intensity of MitoTracker RedCMXRos, was significantly reduced by acute glucose deprivation but restored by repeated episodes of glucose deprivation (Fig 8B).

To evaluate mitochondrial function further, we performed bioenergetic profiling in cells previously treated with single or repeated episodes of glucose deprivation and then assessed at 10 mM glucose to determine if repeated low glucose treatment altered ATP production. Oxygen consumption rate (OCR) and extracellular acidification rate (ECAR) (Fig 8C-E) progressively declined from control levels with one or 3 previous episodes of glucose deprivation but in cells treated with 5 previous episodes of glucose deprivation, both OCR and ECAR were at levels observed in cells without previous glucose deprivation. Hypoglycemia did not affect non-mitochondrial oxygen consumption, as seen by comparable OCR amongst the treatment groups following inhibition with rotenone and antimycin A (Fig 8C). Acute hypoglycemia (1 or 3 episodes) progressively decreased ATP production but was partially reversed by 5 episodes of low glucose (Fig 8F). Similarly, both maximal respiration and spare respiratory capacity, calculated as the difference between maximal and basal OCR, were progressively decreased by 1 and 3 episodes and partially compensated with 5 episodes of glucose deprivation (Fig 8G and H). Taken together, these data suggest that repeated glucose deprivation may lead to partial reversal of low glucose-induced changes in ROS and ATP and so impair hypothalamic glucose sensing.

As mitochondrial function and form are closely related, we next examined the effects of acute or repeated glucose deprivation on mitochondrial structure (Fig 9A). Acute glucose deprivation (1x or 3x) increased mitochondrial length by 50% but this effect was absent after 5 episodes of glucose deprivation (Fig 9B). Dynamin related protein 1 (DRP1), a critical regulator of mitochondrial fission, is controlled by post-translational modification. Phosphorylation of DRP (pDRP1) at Ser616 promotes mitochondrial fission while phosphorylation at Ser637 inhibits mitochondrial fission. Acute glucose deprivation (1x or 3x) significantly decreased the intensity of pDRP1(Ser616) immunostaining but further glucose deprivation (5x) blunted this response (Fig 9C). Acute glucose deprivation did not significantly alter pDRP(Ser637) intensity but repeated glucose deprivation (5x) significantly reduced pDRP(Ser637) intensity compared to acute glucose deprivation (Fig 9D). Therefore acute low glucose shifts the balance of mitochondrial fission and fusion leading to elongated mitochondria, which is absent after multiple episodes of low glucose.

**FIGURE 9:**
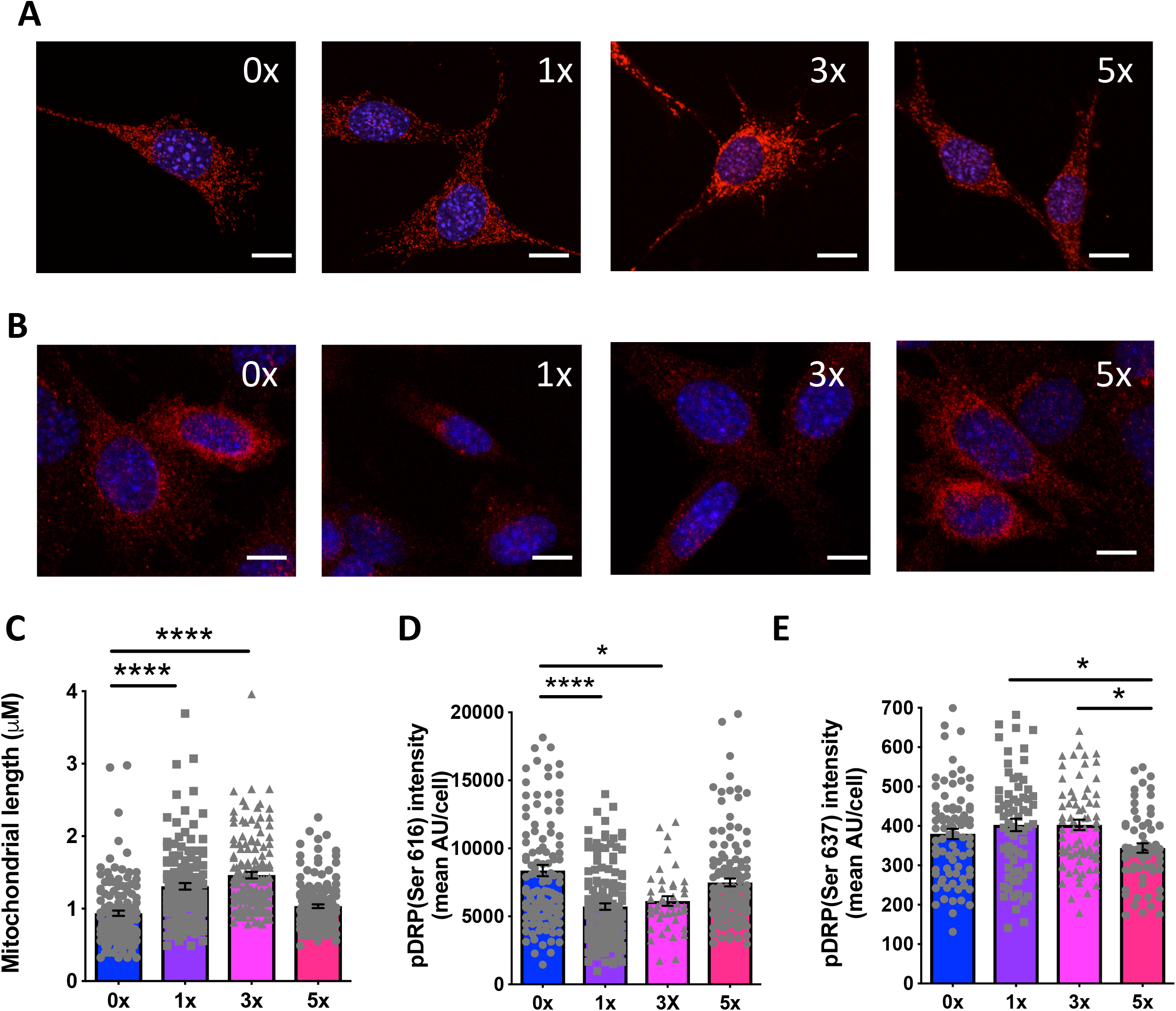
Repeated glucose deprivation blunts effects of low glucose on mitochondrial structure. A) Confocal analyses of Hsp60 immunostaining of mitochondria in N38 cells after none (0x), single (1x) or repeated glucose deprivation (3x and 5x). Scale bar: 10 µm B) Confocal analyses of pDRP Ser616 immunostaining of mitochondria in N38 cells after none (0x), single (1x) or repeated glucose deprivation (3x and 5x). Scale bar: 10 µm C) Quantification of mitochondrial length in N38 cells after none (0x), single (1x) or repeated glucose deprivation (3x and 5x; n = 4 experiments, 140 – 160 mitochondria/group).**** p < 0.0001, χ^2^_(3)_ = 125.4, Kruskal-Wallis test with Dunn’s multiple comparison test. D) Quantification of pDRP(Ser 616) intensity in N38 cells after none (0x), single (1x) or repeated glucose deprivation (3x and 5x; n = 4 experiments, 40 – 130 cells/group). * p = 0.03, **** p < 0.0001, χ^2^_(3)_ = 58.7, Kruskal-Wallis test with Dunn’s multiple comparison test. E) Quantification of pDRP(Ser 637) intensity in N38 cells after none (0x), single (1x) or repeated glucose deprivation (3x and 5x; n= 4 experiments, 63 – 81 cells/group). * p = 0.01 vs. 1x and vs. 3x, F_(3, 280)_ = 3.875, one-way ANOVA with Tukey’s multiple comparison test. Data are displayed as mean ± SEM.

### Mdivi-1 preserves the response to low glucose after repeated glucose deprivation

Our data indicate that acute low glucose leads to mitochondrial fusion in GHRH neurons but repeated hypoglycemia increases mitochondrial fission and reduces neural activity. To address whether maintaining mitochondrial structure could preserve sensitivity to low glucose in cells treated with multiple episodes of low glucose, we used the small molecule DRP1 inhibitor, mitochondrial division inhibitor-1 (mdivi-1). Mdivi-1 inhibits mitochondrial fission in a diverse range of mammalian cells (28). Mdivi-1 maintained mitochondrial length (Fig 10A) and preserved low glucose induced depolarization in N38 cells treated with repeated glucose deprivation (Fig 10B). In cells without previous glucose deprivation, there was a non-significant decrease in the voltage response to glucose deprivation. These findings suggest mdivi-1 treatment prevents mitochondrial fission and maintains responsiveness to hypoglycemia despite repeated previous episodes of hypoglycemia. To confirm these findings *in vivo*, we examined the effects of mdivi-1 on the glucose and c-fos response to repeated glucopenia. Combined daily treatment with 2DG and mdivi-1 partially reversed the blunted glucose response to repeated glucopenia (Fig 10C) and restored GHRH neuron activation (Fig 10D). These findings suggest inhibiting mitochondrial fission *in vivo* maintained GHRH neuron sensitivity to glucopenia even after repeated glucose deprivation.

**FIGURE 10:**
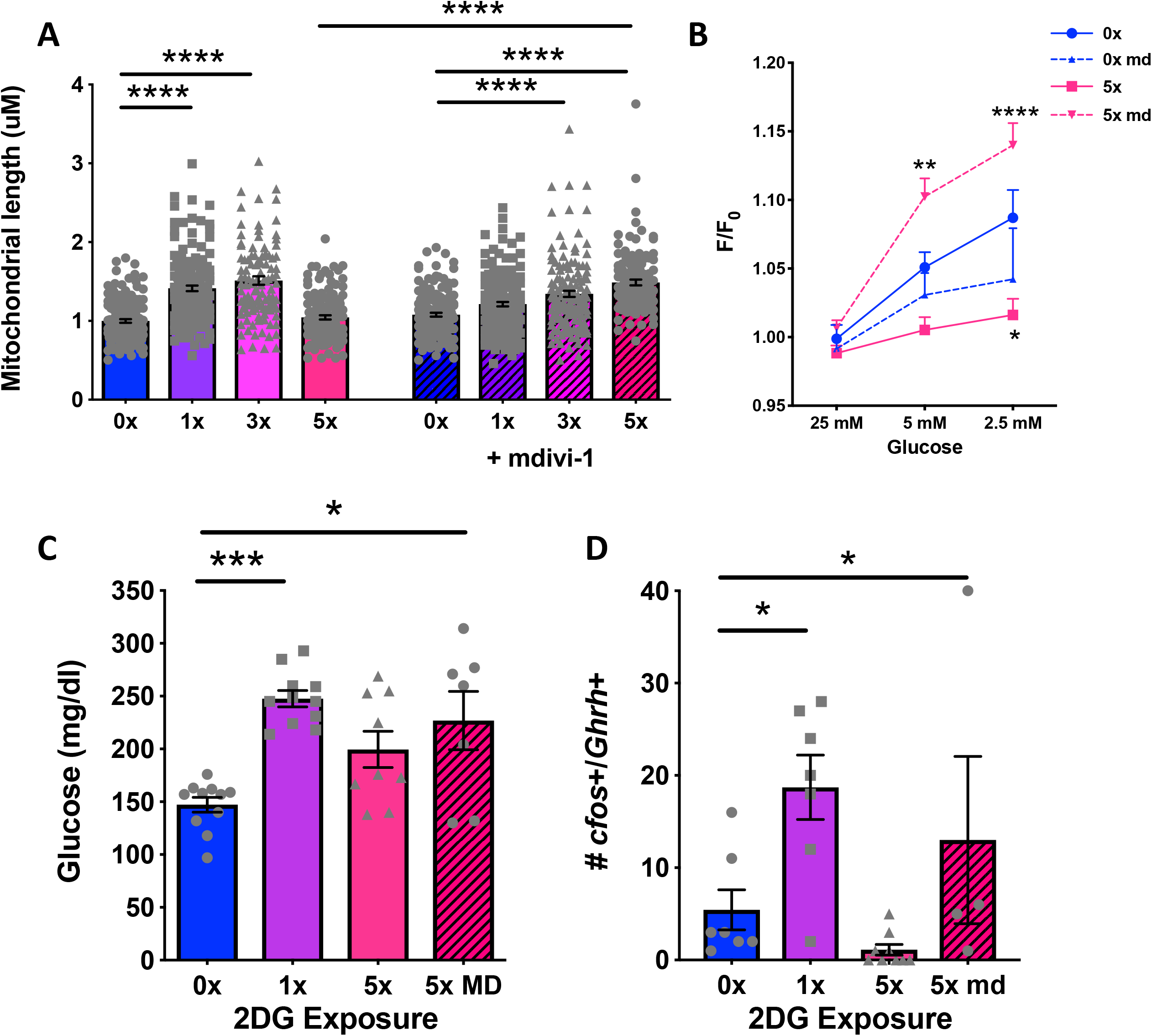
Mdivi-1 preserves the response to low glucose after repeated glucose deprivation. A) Quantification of mitochondrial length in N38 cells after none (0x), single (1x) or repeated glucose deprivation (3x and 5x; n = 3 experiments, 110 – 168 mitochondria/group) with and without mdivi-1 treatment. **** p < 0.0001, F_(7, 1082)_ = 35.33, one-way ANOVA with Tukey’s multiple comparison test. B) Quantification of peak fluorescence (F/F_0_) with 25 mM, 5 mM and 2.5 mM glucose treatment in N38 cells with and without previous glucose deprivation, with and without mdivi-1 treatment. **** p < 0.0001 5x with mdivi-1 vs. 5x at 2.5 mM glucose, ** p = 0.003 5x with mdivi-1 vs. 5x at 5 mM glucose, * p = 0.01 5x vs. 0x at 2.5 mM glucose. Significant effect of mdivi-1 treatment p < 0.0001, F_(3, 1482)_ = 9.187, two-way ANOVA with Tukey’s multiple comparison test, n = 68 – 147 cells/group. C) Analysis of blood glucose after vehicle (0x), single (1x), repeated (5x) i.p. 2DG administration or repeated 2DG administration with mdivi-1 treatment (5X MD). *** p = 0.0004, * p = 0.02, χ^2^_(3)_ = 15.67, Kruskal-Wallis test with Dunn’s multiple comparison test, n = 7 – 11/group. D) Quantification of *c-fos*-/*Ghrh*-positive cells after vehicle (0x), single (1x) or repeated (5x) i.p. 2DG administration or repeated 2DG administration with mdivi-1 treatment (5x MD). * p = 0.04 0x vs. 1x, * p = 0.01 0x vs. 5x MD, Welch’s F_(3, 7.869)_ = 8.338, Welch ANOVA test with Dunnett’s multiple comparison test, n = 4 – 9/group. Each dot represents data from individual cells or animals and data are displayed as mean ± S.E.M.

## Discussion

Our studies demonstrate GHRH neurons are a distinct ARC population of multipolar neurons with polysynaptic connections to the pancreas that are activated by acute glucopenia *in vivo*. Repeated glucopenia leads to substantial remodeling of excitatory and inhibitory inputs to reduce activation of GHRH neurons. This is accompanied by evidence of microglial activation. Using a novel *in vitro* model to reproduce the blunted response to glucose deprivation, we demonstrate that repeated low glucose impairs neuropeptide release, intracellular calcium and hyperpolarizes GI neurons. Repeated glucose deprivation results in partial restoration of ATP production and mitochondrial fission. Pharmacological inhibition of mitochondrial fission maintains responses to low glucose despite preceding glucopenia both *in vitro* and *in vivo*. These studies demonstrate that repeated glucopenia leads to structural and functional remodeling of hypothalamic circuits and GI neurons attributable, in part, to increased mitochondrial fission. Inhibitors of excess mitochondrial fission, such as DRP1 inhibitors, may be effective strategies to prevent hypoglycemia unawareness.

The CRR to hypoglycemia is multifaceted with behavioral, neural and hormonal components. Our studies indicate GHRH neurons may contribute to several aspects of CRR. A critical behavioral response to low glucose is increased food intake and recent reports show non-AGRP, non-POMC arcuate neurons play a role in feeding (29). Our data demonstrate that GHRH neurons are GABAergic and form a distinct population that do not overlap with AGRP or POMC expression. Therefore GHRH neurons could contribute to hypoglycemia induced feeding. In addition, we find that GHRH neurons are synaptically connected to the pancreas so hypoglycemia-activated GHRH neurons could directly influence pancreatic endocrine function via neural pathways. Further work examining the feeding and hormonal responses to targeted GHRH neuron activation and silencing will be needed to determine their roles in hypoglycemia-induced feeding, suppression of insulin and increased glucagon release.

The hormonal responses to hypoglycemia include increased ACTH and GH and indeed, insulin-induced hypoglycemia is the gold-standard test for somatotrophic axis function in clinical practice (30). However, the mechanisms by which hypoglycemia increases GH are not fully understood. In clinical studies, pretreatment with GHRH to induce insensitivity to GHRH release does not prevent GH release with hypoglycemia suggesting insulin-induced hypoglycemia can increase GH independent of GHRH release (31). There are also species differences in the GH response to hypoglycemia. In rats, hypoglycemia reduces plasma GH (32). In sheep, the GH response depends on nutritional state and insulin dose. Fasted sheep increase GH in response to hypoglycemia with late suppression at higher insulin doses and no increase in fed sheep (33). In mice, hypoglycemia has been reported to decrease (34) or have no effect (35) on GH. Here, we found acute glucose deprivation in 5-hour fasted mice reduced plasma GH at 90 minutes. Pituitary GH release is regulated by numerous factors in addition to GHRH. For example, SST suppresses GH release, can over-ride GHRH and is reported to be elevated with low glucose and 2DG in rats (36). In addition, GABA, which colocalizes with GHRH in our studies, inhibits GH release (37). However, it is possible that GH suppression may be an appropriate CRR. In some studies, GH showed a biphasic action on glucose metabolism with a transient stimulation of glucose uptake followed by a more prolonged inhibition of glucose uptake and increased blood glucose(38). Longitudinal studies of GH release with hypoglycemia may provide additional insight into this aspect of counter-regulation in mice.

Repeated glucopenia led to a substantial shift in the balance of excitatory and inhibitory inputs onto GHRH neurons. Excitatory dendritic spines were absent and somatostatin terminals, a major inhibitory input on GHRH neurons, were markedly increased. These modified inputs to GHRH neurons were accompanied by significant microglial activation, known to be involved in synaptic remodeling (39). Detailed electrophysiological studies to assess excitatory and inhibitory postsynaptic currents in response to acute and repeated glucopenia would confirm the shift to increased inhibitory inputs on GHRH neurons. Similar alterations in inputs have been reported in ARC AGRP neurons in response to nutritional status (40) and there is synaptic remodeling in VMH neurons with repeated hypoglycemia (41). While GHRH neurons are known to have both glutamatergic and GABAergic inputs, the sites and neurochemical identity of the neural populations that provide these inputs are unknown. Monosynaptic retrograde tracing from other ARC neural populations identified inputs from multiple hypothalamic, midbrain and other brainstem regions (42) and is in keeping with the multipolar morphology of GHRH neurons that is typical of integrative neurons. Mapping the sites and neurochemical identity of GHRH inputs would be an initial step in determining how GHRH neurons contribute to glucose sensing.

In keeping with our previous *ex vivo* studies (10), GHRH neurons are activated by low glucose *in vivo.* This is likely via cell autonomous glucose sensing as there were no significant changes in dendritic spines or SST terminals with acute glucopenia, although we cannot exclude alterations in the strength of excitatory inputs that occur without dendritic spine remodeling or in non-SST-expressing inhibitory inputs. The GI GHRH-expressing hypothalamic cell line, N38, recapitulated many of the *in vivo* responses in GHRH neurons including activation and increased GHRH with low glucose and a blunted calcium and GHRH response to repeated low glucose. Our findings are similar to those observed in VMH GI neurons, with low glucose-induced activation that is blunted by previous hypoglycemia (21). Calcium influx in hypothalamic GI neurons is via voltage-gated calcium channels and blunted calcium influx could occur through depolarization block. This is a proposed mechanism for impaired glucagon release with hyperglycemia (43). Our voltage imaging studies suggest that repeated low glucose leads to hyperpolarization of GHRH neurons, rather than depolarization block. Previous work has implicated ATP-sensitive chloride channels such as the cystic fibrosis transmembrane receptor (CFTR) in silencing of GI neurons with increased glucose (44),(45). Increased intracellular ATP, which we observe with repeated low glucose, would open CFTR to increase chloride currents, leading to hyperpolarization even when glucose is low.

Mitochondrial dynamics are critical to glucose sensing in ARC and VMH GE neurons (13). The effects of nutritional signals are cell type-specific. In VMH GE neurons, high glucose reduces ROS and mitochondrial length. Loss of mitochondrial remodeling in these neurons prevents glucose sensing leading to systemic glucose abnormalities (46). Similarly in ARC GE neurons, genetic disruption of mitochondrial remodeling alters ARC glucose sensing resulting in disrupted glucose metabolism (14). We have extended these findings by demonstrating that mitochondrial dynamics contribute to glucose sensing in GI neurons. Similar to previous studies that show a role for disrupted mitochondrial dynamics in abnormal glucose metabolism secondary to high-fat diet, our data support the concept that mitochondrial abnormalities contribute to the blunted glucose response with repeated glucopenia. Together, this suggests that abnormal mitochondrial structure may be a common mechanism leading to reduced glucose sensing with either prolonged hyper- or hypoglycemia. Additional factors are likely to regulate mitochondrial dynamics in response to low glucose. For example, uncoupling protein 2 (UCP2) has been implicated in regulation of mitochondrial fission and fusion in ARC and VMH neurons with high glucose (46, 47). Further studies are needed to identify the role of UCP2 or other pathways in mitochondrial remodeling with hypoglycemia.

Targeting mitochondrial fission may provide a strategy to prevent hypoglycemia unawareness. In VMH and ARC GE neurons, genetic restoration of mitochondrial remodeling restores glucose sensing and normalizes blood glucose (14, 46). Here we chose to use a pharmacological approach to reduce mitochondrial fission that might have translational applications. The small molecule DRP1 inhibitor, mdivi-1, preserved mitochondrial length with repeated low glucose and maintained responsiveness to low glucose even after repeated episodes *in vitro* and *in vivo*. Our *in vitro* findings suggest mdivi-1 improves glucose sensing through intracellular effects but it may also restore the balance of excitatory and/or inhibitory inputs onto GHRH neurons. Systemic mdivi-1 has been reported to improve synaptic function in neurodegenerative diseases such as Alzheimer’s and Parkinson’s diseases. Mdivi-1 has been used in multiple preclinical studies and found to have beneficial effects, particularly on neuronal function (48). However, detailed assessments are needed to determine its exact mechanism. Mdivi-1 inhibits mitochondrial fission via actions on DRP1 but DRP1 plays roles in multiple downstream pathways (49). Mdivi-1 has also been reported to have DRP1-independent effects on mitochondrial complex I to attenuate ROS production (50). Further studies are required to understand and selectively target the DRP-1 dependent and independent pathways involved to determine their contributions to preventing hypoglycemia unawareness.

In conclusion, GHRH neurons form a distinct population of GABAergic ARC neurons that are synaptically connected to the pancreas. These neurons are activated by low glucose *in vivo*. Repeated glucose deprivation blunts GHRH neuron activation through circuit-wide and cell intrinsic effects. Excitatory inputs are diminished and inhibitory inputs upregulated with marked microglial activation *in vivo*. Using a new *in vitro* model of blunted glucose sensing, we show repeated glucose deprivation reduces GHRH release, diminishes calcium influx and hyperpolarizes GHRH neurons. Repeated glucose deprivation blunts the effects of low glucose on ATP production and mitochondrial elongation. Maintaining mitochondrial length by pharmacological inhibition of mitochondrial fission preserves responsiveness to hypoglycemia *in vitro* and *in vivo* even after repeated glucopenia. Together these findings suggest structural and functional remodeling of hypothalamic circuits and glucose-sensing neurons contribute to decreased responsiveness to low glucose. Approaches that target mitochondrial dynamics may offer new approaches to prevent hypoglycemia unawareness.

## Methods

### Animals

Male and female C57BL6J and GHRH-GFP heterozygous mice (C57BL6J background) (12 – 30 weeks), randomized by body weight, were housed under controlled light conditions (12 h light/12 h dark) and temperature (22°C) and fed *ad libitum* on standard mouse chow. GHRH-GFP mice were a gift from Dr. Iain Robinson (MRC National Institute for Medical Research, London, UK). Animal care and experimental procedures were performed with the approval of the Animal Care and Use Committee of Icahn School of Medicine at Mount Sinai under established guidelines.

### In vivo studies

Animals were randomized by body weight. Animals were fasted (5 h) in their home cage before treatment with intraperitoneal (i.p.) saline or 2-deoxy-D-glucose (Sigma, 2DG, 400 mg/kg). Animals were treated on five consecutive days receiving saline alone (0x), saline for 4 days and 2DG on day 5 (single glucose deprivation, 1x) or 2DG for 5 days (repeated glucose deprivation, 5x). To assess the effects of inhibition of mitochondrial fission *in vivo*, mice were treated as described above with injection of mdivi-1 (Sigma, 40 mg/kg in corn oil) or vehicle at the same time as saline or 2DG on each treatment day. Glucose readings were taken from tail blood at 90 min. For *in situ* hybridization studies, mice were sacrificed 30 mins after final injection. For measurement of counter-regulatory hormones, retro-orbital blood was taken 90 mins after injection. Blood samples were centrifuged, and plasma stored at −80°C until analysis began.

For retrograde tracing studies, GHRH-GFP mice were induced and maintained on isoflurane anesthesia (2%) before injection of pseudorabies virus expressing red fluorescent protein (PRV-RFP) (5 x 100 nl) into the epigonadal fat pad, pancreas, gastrocnemius muscle, adrenals or liver (n = 3-5 per organ). After viral injection the needle was left in place for 10 min. The skin was closed with stainless steel clips and appropriate analgesia given. Five to seven days later, mice were perfused with 4% paraformaldehyde (PFA) and brains removed for immunohistochemical analysis.

### Assays

Blood glucose was determined using a Breeze 2 glucometer (Bayer; Leverkusen, Germany). Plasma levels of counter-regulatory hormones and GHRH levels in cell supernatant were determined by ELISA (Crystal Chem Glucagon 81518; Corticosterone 80556; GH Millipore Sigma 3P EZRMGH45K, GHRH Aviva systems biology OKEH03121).

### Iontophoretic dye injection, neuronal imaging, 3-dimensional neuron reconstruction, and morphological analysis

GHRH-GFP mice (6-12 weeks old) were treated as described above (n = 4-8). At 90 mins after the final treatment, mice were perfused transcardially with phosphate-buffered saline (PBS) then 1% paraformaldehyde (PFA) for 2 min, followed by 4% PFA/0.125% glutaraldehyde in phosphate buffer of neutral pH for 10 min. Brains were removed and postfixed in the same fixative for 6 hours, then stored in PBS. Coronal sections (125 µm-thick) were cut on a vibratome (Leica VT1000S) and mounted on a nitrocellulose membrane filter in phosphate buffer. Using an epifluorescence microscope (Leica DM LFS) equipped with a micromanipulator and a current generator, GHRH-GFP expressing neurons were identified for iontophoretic dye injection under a 40X objective. Once identified, a glass pipette filled with 5% Alexa 555 dye was used to penetrate the neuronal cell body and a direct current (5-10 nA) was applied to ensure filling of the entire cell. About 1 to 3 cells were injected on each section, spaced so as to prevent overlap of their dendrites. Tissue sections with filled cells were mounted onto SuperFrost slides using Fluoromount G and coverslipped using spacers.

Images of dendritic spines were captured and analyzed as previously described (51). The investigator was blinded to treatment group. Briefly, Z-stacks of whole cells were acquired at a resolution of 1024 x 1024 pixels, at a Z-step of 0.1 µm, and optimized for gain and offset, using a Zeiss LSM780 confocal microscope with a 100x/1.4 N.A. Plan-Apochromat objective, a digital zoom of 3x and an Ar/Kr laser at an excitation wavelength of 550 nm. In total, dendritic segments from 29 neurons were examined, with an average of 2 neurons each from 16 animals, yielding 656 µm of dendritic processes were quantified. Confocal images were deconvolved using an iterative blind deconvolution algorithm (AutoQuant X3) and analyzed manually for spine density and classification after volumetric reconstruction using Imaris software (Bitplane, Zurich, Switzerland).

Images of dendritic spines were captured and analyzed as previously described (51), from dendritic segments selected at 0 – 200 µm from the soma. The selection criteria for dendritic spines is as previously described (52). Briefly, Z-stacks of dendritic segments were acquired at a resolution of 1024 x 1024 pixels, at a Z-step of 0.1 µm, and optimized for gain and offset, using a Zeiss LSM780 confocal microscope with a 100x/1.4 N.A. Plan-Apochromat objective, a digital zoom of 3x and an Ar/Kr laser at an excitation wavelength of 555 nm. In total 29 neurons were examined, with an average of 2 neurons from 16 animals, yielding 656 µm of dendritic processes. Confocal images were deconvolved using an iterative blind deconvolution algorithm (AutoQuant X3) and analyzed manually for spine density and classification after volumetric reconstruction using the Imaris software. Spines were classified into filopodia/thin, stubby, and mushroom based on the head to neck diameter ratio and maximum head diameter.

### iDISCO+ brain clearing

GHRH-GFP mice (n = 3) were anesthetized with isofluorane then perfused with PBS followed by 4% PFA and brains then post-fixed in PFA at 4°C overnight. iDISCO+ was performed as described previously (Renier et al., 2016). In brief, brains had their hindbrains and olfactory bulbs removed to reduce sample sizes, and were dehydrated with a methanol gradient (20%-100%) at room temperature (RT), incubated in 100% dichloromethane (DCM) (Sigma-Aldrich, St. Louis, MO, USA) to remove hydrophobic lipids, and bleached in 5% H_2_O_2_ overnight at 4°C to reduce tissue autofluorescence. Brains were then rehydrated with a methanol gradient (80%-20%) and incubated in permeabilization buffer with 5% DMSO/0.3 M Glycine in 0.1% Triton X-100/0.05% Tween-20/0.0002% heparin/0.02% NaN_3_ (PTxwH) in 1x PBS for 1 day. Brains were incubated in blocking buffer consisting of PTxwH with 3% normal donkey serum (Jackson Immunoresearch, West Grove, PA, USA) at 37°C overnight. Immunolabeling for GFP (Aves, Table 1) was performed in blocking buffer for 4 days at 37°C, and was followed by 5 washes in PtxwH and secondary antibody incubation for 4 days at 37°C. Samples were washed (5 times each) in PtxwH at 37°C and in PBS at RT. Optical clearing was achieved by dehydrating with a methanol gradient (20%-100%), washing 3 times for 30 min in 100% methanol followed by 3 times for 30 min in DCM, before transferring the samples to dibenzylether (DBE) (Sigma-Aldrich). The hypothalamus was imaged with an Ultramicroscope II (LaVision BioTec, Bielefeld, Germany) using a 4x objective, withZ-stacks acquired at a Z-step of 3 µm giving an average of 492 sections per animal. Hypothalamic images were analyzed with Imaris for quantification of GFP-positive neurons using the Imaris “spot” detection algorithm (see next section). The count was not corrected to the brain volume.

**Table 1:** Primary antibodies used. Summary of primary antibodies used in the studies with source, catalog number, host species and dilution.

| Antigen | Company | Catalog number | Host species | Dilution | Incubation |
| --- | --- | --- | --- | --- | --- |
| Somatostatin (54) | Immunostar | R210-01 | Rabbit | 1:200 | 48 Hours |
| GFP (22) | Aves | GFP-1020 | Chicken | 1:1000 | Overnight |
| IBA-1 (55) | Wako | 019-19741 | Rabbit | 1:1000 | Overnight |
| pDRP s616 (56) | Cell Signaling | 4494S | Rabbit | 1:200 | 72 Hours |
| pDRP s637 (57) | Cell Signaling | 4867S | Rabbit | 1:1000 | Overnight |
| HSP60 (58) | Santa Cruz | sc-271215 | Mouse | 1:500 | Overnight |
| RFP (59) | Rockland | 600-401-379 | Rabbit | 1:1000 | Overnight |
| CFOS (59) | Cell Signaling | 2250S | Rabbit | 1:1000 | Overnight |
| AGRP (10) | Phoenix | H-003-57 | Rabbit | 1:500 | Overnight |
| NPY (10) | Phoenix | H-049-03 | Rabbit | 1:1000 | Overnight |
| POMC (10) | Phoenix | H-029-30 | Rabbit | 1:1000 | Overnight |

### Immunocytochemistry, immunohistochemistry and in situ hybridization

Immunocytochemistry (ICC) and immunohistochemistry (IHC) were used to detect expression of heat-shock protein 60 (HSP60), phosphoDRP Ser616, phosphoDRP Ser 637 in cells, GFP, RFP, POMC, NPY, AGRP, SST, and IBA1 in tissue. For ICC, cells grown on fibronectin-coated coverslips were washed twice in PBS and then fixed for 15 min in 4% PFA (Electron Microscopy Services, Hatfield, PA). For IHC, isoflurane anesthetized mice were transcardially perfused with 0.9% saline followed by 4% PFA in 0.1 M phosphate buffer. Brains were removed, immersed in 4% PFA at 4°C overnight. Coronal slices were cut at 50-µm thickness on a vibratome (Leica VT1000). Immunofluorescence and double labeling were performed as previously described (10). Briefly, cells or tissue was washed in 0.03 M Triton-X 100 (TX) in PBS followed by incubation in blocking solution (1 h, 3% bovine serum albumin in 0.03 M TX/0.1 M PBS). Cells or tissue were incubated in primary antibody in blocking solution (16-72 hours depending on target protein, 4°C; see Table 1 for details). Cells or tissue were rinsed in TX/PBS and incubated in blocking solution containing secondary antibody (Alexa conjugated, Life Technologies, 1:1000, 2 hours, RT). After staining, cells or tissue sections were mounted using Fluoromount. For IHC to detect NPY, POMC and AGRP, GHRH-GFP mice were treated with colchicine to enhance cell body staining, as previously described (10). Perfusion, sectioning and processing were performed as described above.

For *in situ* hybridization, brains were snap-frozen in liquid nitrogen and 30-µm thick sections were cut on a cryostat. An RNAscope multiplex fluorescent assay kit was used (Advanced Cell Diagnostics, Newark, CA; catalog #323100). Briefly, sections were fixed in 4% PFA for 1 hr at 4°C then dehydrated in ethanol followed by protease III treatment for 30 min. Sections were hybridized with RNAscope oligonucleotide probes against *Ghrh* (#470991), *Gck* (#400971), *Sst* (#404631), *fos* (#316921), *Gad2* (#439371), and/or *Slc17a6* (#319171) then incubated with signal amplifier and TSA Plus fluorophores (PerkinElmer Inc., Waltham, MA; catalog # NEL744001KT, NEL745001KT, NEL741001KT) according to RNAScope protocols. Slides were washed with RNAscope wash buffer, counterstained with DAPI and mounted with Prolong Gold mounting medium.

Images were acquired as z stacks using a Zeiss Inverted LSM 780 laser scanning confocal microscope (Carl Zeiss MicroImaging, Inc.). Single optical sections were collected with the pinhole set to 1 Airy Unit for the far red channel and adjusted to the same optical slice thickness for the remaining channels. Bleed-through was assessed by examining single-labeled samples for emission in the appropriate laser detection channel when the corresponding laser was deactivated and none was observed. Briefly, Z-stacks were acquired at a resolution of 1024 x 1024 pixels, at a Z-step of 1 µm, and optimized for gain and offset, using a Zeiss LSM780 confocal microscope with a 20x/1.4 N.A. Plan-Apochromat objective, a digital zoom of 1 and an Ar/Kr laser at an excitation wavelengths of 358 nm, 499 nm, 591 nm. For the quantification studies, exposure times were set based on control group and were comparable among the different experimental groups. Staining intensity, number of cells and colocalization were quantified using the Imaris software and profile-based counting method. The investigator was blinded to treatment group. Briefly, for immunohistochemistry in brains from GHRH-GFP, every 3^rd^ section was collected, immunostained and imaged. For *in situ* hybridization, every 6^th^ section was collected, probed and imaged. Confocal image stacks were reconstructed and visualized as 3-dimensional volumes with Imaris software. The morphological limits of the ARC were recognized using DAPI nuclear staining and regions of interest (ROI) were drawn around the ARC. The Imaris “spot” detection algorithm was used for semiautomatic identification, counting of fluorescently labeled cells, colocalization of cells and of terminals close to GHRH neurons. The parameters used were an object size of 8 µm diameter for cells and of 0.4 µm for SST-immunoreactive terminals, with background subtraction and a threshold of 32 for GHRH neurons. We considered spots that were within 0.5 µm to be colocalized. To quantify gene expression signal, the area and intensity of > 20 dots were measured to determine the mean intensity per dot. ROI were drawn around each *Ghrh* neuron and the total signal area and intensity in each ROI measured then normalized for the mean intensity per dot to give the gene signal within each *Ghrh* neuron.

### Quantitative real-time PCR

Total RNA was extracted from N38 cells using RNAeasy plus kit (Qiagen, MD) according to the manufacturer’s protocol. RNA purity and quantity was assessed by spectrophotometry (Spectramax, Molecular Devices) and RT-PCR was performed using an ABI 7500 system as previously described (10) using the following primers: GHRH: Primer 1 CTGTATGCCCGGAAAGTGAT, Primer 2 – GTCATCTGCTTGTCCTCTGTC, Probe – TCCTCTCCC. RPL23, Primer 1 - TTAGCTCCTGTGTTGTCTGC, Primer 2 - ACTTCCTTTCTGACCCTTTCC, Probe: TTCGACATCTTGAACG.

### Cell culture and in vitro studies

Murine hypothalamic neural cells (N38, Cellutions Biosystems, *Mycoplasma* testing and STR profiling performed by Cellutions Biosystems) were grown in normal culture medium comprised of high glucose (25 mM) Dulbecco’s modified eagle medium (DMEM)/10% fetal bovine serum (FBS) at 37°C and 5% CO_2_. For immunocytochemistry studies, cells were grown on fibronectin coated 12-mm cover glass (Fisher Scientific, Pittsburgh, PA).

For acute or repeated glucose deprivation, culture medium was replaced with no glucose DMEM/10% fetal bovine serum supplemented with 25 mM (control) or 2.5 mM glucose (glucose deprivation) for 2 h. Normal culture medium was replaced at the end of this period. Cells were treated with no glucose deprivation (0x, 5 treatments with 25 mM glucose/DMEM/10% FBS), acute glucose deprivation (1x, 4 treatments with 25mM glucose/DMEM/10% FBS and 1 treatment with 2.5 mM glucose/DMEM/10% FBS), repeated glucose deprivation (3x), 2 treatments with 25 mM glucose/DMEM/10% FBS and 3 treatments with 2.5 mM glucose/DMEM/10% FBS or 5x, 5 treatments with 2.5 mM glucose/DMEM/10% FBS). To assess the effects of inhibiting mitochondrial fission, N38 cells were treated as described above with the addition of mdivi-1 (50 nM) or DMSO (control). At the end of the final treatment period, either supernatant was removed and stored at - 80°C for GHRH assay, cells were fixed for immunocytochemistry or cells were lysed in RNA lysis buffer (Qiagen, MD) and the lysate stored at −80°C until RNA purification. Each study was repeated on at least 3 occasions with 4 replicates. For examination of mitochondrial membrane potential, N38 cells were incubated with Mitotracker Red CMXRos (100 nM, Invitrogen) in culture medium for the final 30 mins of the last treatment period. Cells were fixed in 4% PFA in culture medium for 15 mins before rinsing in PBS and mounting (Fluoromount with DAPI). Images were acquired as z stacks using a Zeiss Inverted LSM 780 laser scanning confocal microscope (Carl Zeiss MicroImaging, Inc.) as described above. Single optical sections were collected with the pinhole set to 1 Airy Unit and adjusted to the same optical slice thickness for the remaining channels. Fluorescence intensity was quantified using Fiji software with regions of interest (ROI) over individual cells.

### Calcium and voltage Imaging

For calcium imaging studies, N38 cells grown on fibronectin-coated coverglass were loaded with Fluo-4 in Tyrode’s solution (140 mM NaCl, 5 mM KCl, 5 mM HEPES, 1 mM NaH_2_PO_4_, 1 mM MgCl_2_, 1.8 mM CaCl_2_, 25 mM glucose) as previously described (53). Cells were imaged in calcium imaging buffer (53) with 25 mM, 5 mM or 2.5 mM glucose using a Zeiss Axio Observer Z1 inverted widefield fluorescent microscope. Images were acquired every 6 seconds for 3 mins at a resolution of 1024 x 1024 pixels with a 10x Plan-Apochromat objective. For voltage imaging studies, N38 cells grown on chamber slides (Ibidi) were washed twice then loaded with Fluovolt (Invitrogen) in Tyrode’s solution according to the manufacturer’s instructions. Imaging was performed in Tyrode’s solution with 25 mM, 5 mM or 2.5 mM glucose in 5% CO_2_ at 37°C. Briefly, images were acquired every second for 180 seconds at a resolution of 512 x 512 pixels and optimized for gain and offset, using a Zeiss LSM780 confocal microscope with a 20x/1.4 N.A. Plan-Apochromat objective, a digital zoom of 1 and an Ar/Kr laser at an excitation wavelength of 495 nm. All imaging studies were performed on three occasions for each condition. Fluorescence intensity was quantified using Fiji software with ROI over individual cells (calcium imaging) or cell membranes (voltage imaging).

### Measurement of oxygen consumption rate

Oxygen consumption rate (OCR), extracellular acidification rate (ECAR), ATP production, maximal respiration and spare respiratory capacity were determined in a Seahorse XFe24 extracellular flux analyzer. Measurements were performed at the end of the final glucose deprivation treatment in medium with 10 mM glucose and 1% FBS. Basal measurements were determined, and OCRs were recorded after the following additions: oligomycin (2.5 µmol/l), carbonyl cyanide 4-(trifluoromethoxy)-phenylhydrazone (FCCP; 1 µmol/l), rotenone (1 µmol/l) and antimycin A (1 µmol/l).

### Measurement of reactive oxygen species

For measurement of ROS, N38 cells were cultured on 96 well plates and treated as described above. For the final hour of treatment, Amplite ROS Green (AAT Bioquest) was added to the culture medium. Fluorescence signal was measured by microplate reader (Spectramax, Molecular Devices).

### Statistics

All the data shown are the mean ± S.E.M of results of a number of experiments (indicated in the figure legends). Distribution was assessed by Kolmogorov-Smirnov test and Shapiro-Wilk test. Significance was determined by the unpaired two-way *t* test, one-way ANOVA with the post hoc Tukey (Gaussian distribution), Welch ANOVA (Gaussian distribution with unequal standard deviation), Kruskal-Wallis test (non-parametric distribution) or two-way ANOVA with Sidak’s multiple comparison test for repeated measures. Significance was set at an alpha level of 0.05.

## Author contributions

MB performed experiments, analyzed data and contributed to the writing of the manuscript. AA, KD, RL, MJG, KC performed experiments and reviewed the manuscript. MV, MNS, JEC, PRH provided experimental and intellectual expertise. SAS performed experiments, analyzed data and wrote the manuscript. MB and SAS designed the studies. All authors discussed the results and edited the manuscript.

## Acknowledgements

A.A. is supported by a postdoctoral fellowship from the Charles H Revson Foundation. K.D. is supported by a predoctoral fellowship from the American Heart Association (18PRE33960254). Supported by the American Diabetes Association Pathway to Stop Diabetes Grant ADA #1-17-ACE-31.This work was supported in part by grants from the National Institutes of Health National Institute of Mental Health (U01MH105941), National Institute of Neurological Disease (R01NS097184) and NIH Common Fund (OT2OD024912), Einstein-Mt. Sinai Diabetes Research Center Pilot and Feasibility award (P30DK020541) and Alexander and Alexandrine Sinsheimer Scholar Award.

